# Mass Enhanced Node Embeddings for Drug Repurposing

**DOI:** 10.1101/2022.06.22.497214

**Authors:** Michail Chatzianastasis, Giannis Nikolentzos, Michalis Vazirgiannis

## Abstract

Graph representation learning has recently emerged as a promising approach to solve pharmacological tasks by modeling biological networks. Among the different tasks, drug repurposing, the task of identifying new uses for approved or investigational drugs, has attracted a lot of attention recently. In this work, we propose a node embedding algorithm for the problem of drug repurposing. The proposed algorithm learns node representations that capture the influence of nodes in the biological network by learning a mass term for each node along with its embedding. We apply the proposed algorithm to a multiscale interactome network and embed its nodes (i. e., proteins, drugs, diseases and biological functions) into a low-dimensional space. We evaluate the generated embeddings in the drug repurposing task. Our experiments show that the proposed approach outperforms the baselines and offers an improvement of 53.33% in average precision over typical walk-based embedding approaches.

## 1 Introduction

Recently, there is a growing interest in representing data from different domains in the form of graphs. Indeed, graphs arise naturally in many application domains such as in chemoinformatics [Kearnes et al.(2016)], in physics [Battaglia et al.(2016)], and in natural language processing [Nikolentzos et al.(2020)]. In many cases, machine learning techniques need to be applied to graph-structured data. For instance, some common applications include predicting the estimated time of arrival in Google Maps [Derrow-Pinion et al.(2021)], recommending friends in social media [Fan et al.(2019)] and predicting the quantum mechanical properties of molecules [Gilmer et al.(2017)], just to name a few. These learning tasks focus on different components of graphs such as nodes, edges or subgraphs. For example, common node- and edge-level tasks include predicting the biological state of a protein in a protein-protein interaction network and discovering the type of the relationships between entities in a knowledge graph, respectively.

Traditionally, biology is one of the richest sources of graph-structured data. Protein-protein interaction networks are perhaps the most representative examples of such graphs. Complex diseases usually disrupt several of the proteins in these networks [Huttlin et al.(2017)]. A drug that needs to reach the proteins disrupted by the disease usually binds a single target protein. Through the interactions between proteins, it can eventually affect those proteins that were disrupted by the disease [Barabási et al.(2011)]. Those interactions can be modeled as a graph. This graph can serve as the main tool one can use to investigate the effects of disease treatments and their benefits, i. e., the efficacy of drugs [Guney et al.(2016)], their side effects [Zitnik et al.(2018)], etc. An example of such a graph is the multiscale interactome network [Ruiz et al.(2021)] which was built to capture the biological principles of effective treatments and which consists of disease-perturbed proteins, drug targets and biological functions.

Given the above network, one can apply machine learning techniques to identify how a drug treats a disease. However, such an approach involves several challenges. Perhaps the most important challenge is how to incorporate information about the structure of the graph and potentially of node and edge attributes in the learning model. For example, in the case of the multiscale interactome network, in order to determine whether a drug can treat a given disease, we need to obtain an informative representation of drugs and their proximities to diseases – that potentially is not fully captured by graph statistics and other handcrafted features extracted from the graph such as Jaccard’s coefficient and the Adamic-Adar index [Liben-Nowell and Kleinberg(2007)]. Recently, a lot of attention has been devoted to the development of algorithms that learn continuous representations of different components of graphs, also known as embeddings. Such representations capture the structural information of the underlying graph. In other words, such a methodology maps elements of the graph into a low-dimensional vector space, while the graph structure is preserved. Note also that most embedding approaches are purely unsupervised. Thus, the generated representations can further be used in any downstream machine learning task, e. g., classification or clustering.

Over the past years, a significant amount of effort has been devoted to node embedding algorithms [Cai et al.(2018)]. However, most existing node embedding algorithms are designed to capture different types of node proximities [Tang et al.(2015)] or are motivated by application-specific considerations [Dani et al.(2021)] Such embeddings might fail to capture the full complexity of the interactions between nodes in highly complex networks such as the multiscale interactome network. In this paper, we propose the Mass-Enhanced Random Walk (MERW) algorithm, a new node embedding algorithm which can capture the influence that a node has on other nodes even if they are far from each other in the considered network. The algorithm is inspired by DeepWalk [Perozzi et al.(2014)] which simulates random walks over the input network and also employs ideas from physics. Specifically, the algorithm learns a mass term for each node which capture the node’s importance in the network. Thus, the algorithm can learn whether specific proteins, biological functions and drugs can significantly influence the rest of the network’s nodes. We empirically evaluate the proposed approach in the task of predicting which drugs can treat a given disease. The proposed method outperforms the baselines, thus suggesting that it captures more accurately the biological functions through which target proteins affect the functions of disease-perturbed proteins.

## 2 Related Work

Early methods in the field of graph representation learning follow a simple walk-based approach, i. e., they simulate random walks over the graph and treat the emerging walks as sentences in some special language. They then capitalize on ideas from natural language processing such as the Skipgram model [Mikolov et al.(2013)] to generate node embeddings. DeepWalk [Perozzi et al.(2014)] and node2vec [Grover and Leskovec(2016)] are typical examples of this family of algorithms. The main difference between the two approaches is that the former simulates simple random walks, while the latter performs biased walks. Another family of approaches consists of algorithms which explicitly preserve different types of node proximities. For instance, LINE [Tang et al.(2015)] optimizes an objective function that preserves the first- and second-order proximities, while GraRep [Cao et al.(2015)] preserves high-order proximities by applying SVD to high-order proximity matrices. Several of the above algorithms can be unified into the matrix factorization framework with closed form solutions [Qiu et al.(2018)]. Other embedding approaches instead of node proximities capture different properties of graphs such as their community structure [Wang et al.(2017)] or even edge semantics [Gao et al.(2019)]. Besides the afore-mentioned embedding approaches, graph autoencoders have also recently emerged as a very useful framework for learning node representations. Most of these models, including the variational graph autoencoder [Kipf and Welling(2016)], consist of a graph neural network encoder and a simple inner product decoder. The graph neural network encoder can be replaced with linear models for interpretability purposes [Salha et al.(2020)]. Graph autoencoders have also been generalized to directed graphs [Salha et al.(2019)]. A detailed review of node embedding algorithms is beyond the scope of this paper; we refer the interested reader to the survey on graph embeddings [Cai et al.(2018)].

With regards to the main task of the paper, i. e., predicting which drugs can treat a given disease, previous approaches have used biased random walks to learn a diffusion profile for each drug and disease, which identifies the proteins and biological functions involved in a given treatment [Ruiz et al.(2021)]. Neural network models have also been employed to uncover disease–disease, disease–gene and disease–pathway associations [Gaudelet et al.(2020)]. Recently, graph neural networks were also employed to repurpose drug compounds for the treatment of different human coronavirus diseases [Sugiyama et al.(2021)].

## 3 Methodology

### 3.1 Preliminaries

Let *G* = (*V, E*) be an undirected graph consisting of a set *V* of vertices and a set *E* of edges between them. We will denote by *n* = |*V*| the number of vertices and by *m* = |*E*| the number of edges. Let 𝒩(*v*) denote the set of neighbors of node *v* ∈ *V*, i. e., 𝒩(*v*) ={*u* : (*v, u*) ∈ *E* }. The degree of a node is equal to the number of edges incident to that node, i. e., for a node *v*, deg(*v*) = |𝒩(*v*)|. A walk in a graph *G* is a sequence of vertices *v*_1_, *v*_2_, …, *v*_*k*+1_ where *v*_*i*_ ∈ *V* and (*v*_*i*_, *v*_*i*+1_) ∈ *E* for 1 ≤ *i* ≤ *k*. The length of the walk is equal to the number of edges in the sequence.

In this paper, our objective is to learn an embedding for each node of a graph such that these embeddings capture as much structural information of the graph as possible, but also the interactions between nodes. Formally, we aim to learn a function *f* : *V* → ℝ^*d*^ that maps nodes to feature representations that can be then utilized for some downstream prediction task such as link prediction or influence prediction. *d* is a hyperparameter specifying the number of dimensions of the generated representations.

### 3.2 Mass-Enhanced Random Walk (MERW)

Following previous studies [Perozzi et al.(2014); Grover and Leskovec(2016)], the proposed approach first simulates a number of random walks over the input graph *G*. Specifically, the algorithm starts a specific number of random walks (*t* walks in total) at each node of the graph *v* ∈ *V*. Given the root node of a walk, for a number of steps (which is equal to the length of the walk to be simulated minus one) the algorithm samples uniformly at random one of the neighbors of the last visited node, i. e., one node from the set 𝒩(*v*) where *v* is the last node visited by the walk. In our experiments, we set the length of the random walks to be fixed and equal to *L*. Thus, we end up with *t*|*V*| random walks in total. These walks form the training set of the algorithm.

Then following previous work [Mikolov et al.(2013)], the proposed algorithm maximizes the co-occurrence probability among the nodes that appear within a window *w* in a random walk. More specifically, a window slides across the walk, and given the representation vector of the center node, the proposed algorithm maximizes the probability of its neighbors in the walk. Each node *v*_*i*_ ∈ *V* is associated with two types of embeddings: (1) one employed when the node serves as the central node of the window (embedding **u**_*i*_); and (2) one employed when the node appears in the context of another node (embedding **v**_*i*_). For a center node *v*_*i*_ and of one of its neighbors (within the window) *v*_*j*_, we model the co-occurrence probability as follows:

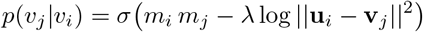

where *λ*∈ *ℝ*^+^ is a hyperparameter, and *m*_*i*_, *m*_*j*_ are trainable parameters (scalar values) which can be thought of as mass parameters. Such mass parameters have been also adopted in previous studies in the context of graph autoencoders [Salha et al.(2019)]. We expect them to capture the importance of the different nodes, i. e., some nodes are more influential than others in the graph. For instance, in a scientific publications citation network, seminal articles are more influential than the rest of the articles. Thus, two influential nodes are more likely to be connected to each other by an edge than two nodes that are not very influential. Regarding the hyperparameter *λ*, it can be tuned by cross-validation. The goal of this parameter is to balance the contribution of the distance of the two node embeddings and of their mass parameters to the loss function. Increasing *λ* forces the model to give more importance to the symmetric node proximity rather than the mass parameters that capture the global influence of a node on its neighbors.

To learn the node embeddings **u**_*i*_, **v**_*i*_ and the mass term *m*_*i*_ for each node *i* ∈ *V*, the proposed model minimizes the following loss function:

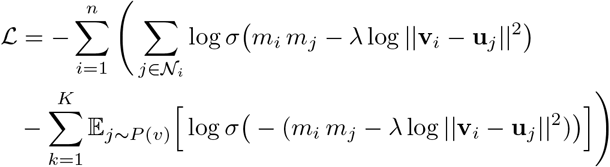

where 𝒩_*i*_ ={ *j* : *u*_*j*_ ∈ 𝒩(*v*_*i*_)}is the set of node indices of the neighbors of node *v*_*i*_, *P* (*v*) is a noise distribution, and *K* is the number of negative samples to be drawn for each node. We optimize the above function using stochastic gradient ascent over the model parameters. The derivatives are estimated using the backpropagation algorithm. The first term of the loss function maximizes the probability of co-occurrence for the central node and nodes that lie in its context window, while the second term randomly samples some nodes that don’t lie in the context window and minimize their probability of co-occurrence. Thus, the objective is to distinguish the nodes that lie in the context window from draws from the noise distribution *P* (*v*) using logistic regression, where *K* negative samples are drawn for each node. The random nodes are sampled based on their frequency of occurrence. Specifically, *P* (*v*) = *U* (*v*)^3/4^*/Z* where *U* (*v*) is a unigram distribution (i. e., probability of finding a specific node in the set of random walks) and *Z* is a normalization constant.

Once the model is trained, we can retrieve the embedding **u**_*i*_ of each node *i* ∈ *V*. The generated representations are general and independent of the downstream prediction task. In this work, we focus on predicting how effective a drug is for a specific disease. Thus, to make predictions, we concatenate the embedding of the drug with that of the disease, and we feed the emerging vector to some standard classifier (e. g., multi-layer perceptron). An illustration of the proposed algorithm is provided in Figure 1.

**Figure 1:**
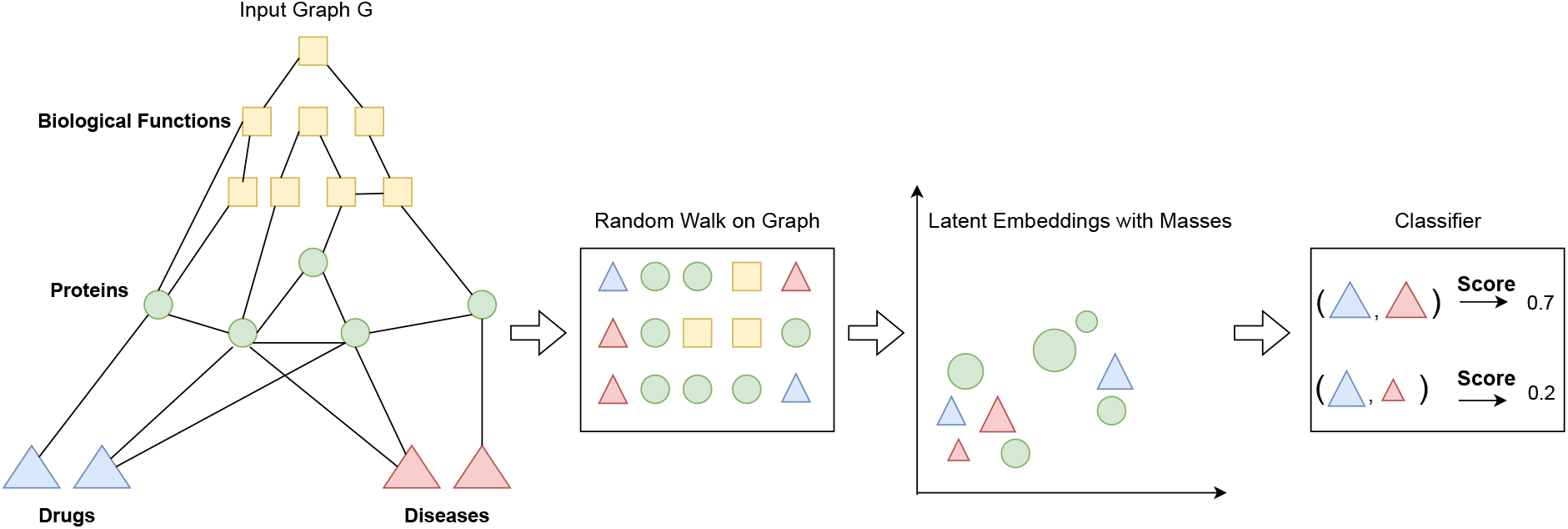
An illustration of the proposed approach. Given a heterogeneous graph *G*, whose nodes correspond to drugs, diseases, proteins and biological functions, the goal is to predict whether a drug can treat a disease. The algorithm first simulates a number of random walks over the graph to produce sequences of nodes. Then, it extracts training samples from those sequences and uses them to learn node embeddings along with mass terms which capture the influence of the nodes in the graph. Finally, pairs of drug and disease embeddings are fed to a classifier to predict the probability of a treatment relationship between them.

## 4 Experiments

Our implementation is publicly available on [github](https://github.com/MichailChatzianastasis/Mass-Enhanced-Node-Embeddings-for-Drug-Repurposing).

### Datasets

We evaluate the proposed node embedding algorithm on a multiscale interactome network [Ruiz et al.(2021)]. It consists of proteins, drugs, diseases and biological functions. Specifically, it uses the physical interaction network between 17, 660 proteins, and the hierarchy of 9, 798 biological functions, in order to capture the principles of effective treatments across 1, 661 drugs and 840 diseases. It contains 478, 728 edges and 5, 926 unique approved drug–disease pairs.

### Experimental setup

We perform 5-fold cross validation. We hold out one of the 5 folds and consider the drugs contained in it as the set of test drugs. The remaining drugs are considered as the train drugs. Subsequently, we create a train set of diseases using all approved indications of the train drugs and a test set of diseases using all approved indications of the test drugs. Within a given split (i. e., “train” or “test”), we evaluate performance using only that split’s sets of drugs and diseases (i. e., for testing performance, we use “test drugs” and “test diseases”). For every disease in the split of interest, each model produces a ranking of the corresponding drugs. To generate the random walks, we set the walk length equal to 10 and the number of walks equal to 20. We use embeddings size equal to 128 and we train the model for 50 epochs. For the classifier we use a multi-layer perceptron with 2 hidden layers, and we train it for 25 epochs.

### Metrics

To evaluate the rankings produced by the different approaches, we use two metrics: (1) Average Precision (AP); and (2) Recall@50. AP is a way to summarize the precision-recall curve into a single value representing the average of all precisions. AP is computed as follows: 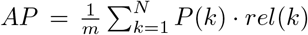, where *m* is the number of relevant items, *N* is the number of recommended drugs, *P* (*k*) is the precision calculated only for the recommendations from rank 1 to *k, rel*(*k*) is an indicator of whether that *k*-th drug was relevant (*rel*(*k*) = 1) or not (*rel*(*k*) = 0). Recall@50 is the proportion of relevant drugs found in the top-50 recommendations.

### Baselines

We compare MERW against the following two node embeddings algorithms: (1) node2vec [Grover and Leskovec(2016)]; and (2) Diffusion Profiles [Ruiz et al.(2021)]. We also compare it against the following methods (we use the results reported in [Ruiz et al.(2021)]): (3) Molecular-scale Interactome; (4) Functional Overlap; and (5) Protein Overlap. The first two baselines use the same input as MERW, but use different approaches to learn node representations. The third baseline uses a molecular-scale interactome as the input graph, which does not exploit the biological functions.

### Results

We report the mean average precision and the mean Recall@50 across the diseases in Table 1. Our model out-performs all the baselines by a large margin. Specifically, MERW offers an increase of 53.33%, 32.89% over node2vec and an increase of 26.37%, 44.38% over Diffusion Profiles in Average Precision and Recall@50 respectively. Our results thus indicate that the proposed algorithm can learn node representations useful for identifying potential drug-disease treatments. We also experiment with different classifiers which all take as input pairs of drug-disease embeddings. The considered classifiers include a multi-layer perceptron (MLP) with 2 hidden layers, a Random Forest classifier (RF) with 100 estimators, and a *k*-nearest neighbors classifier (*k*-NN) with *k* ∈{1, 3, 5}. We observe that MLP achieves the best results, due to its expressive power and its ability to process high dimensional embeddings.

**Table 1:** Reported values averaged across five-fold cross validation on drug-disease treatments dataset [Ruiz et al.(2021)].

| Method | Avg Prec | Rec@50 |
| --- | --- | --- |
| Protein Overlap | 0.064 | 0.298 |
| Functional Overlap | 0.050 | 0.237 |
| Molecular-scale Interactome | 0.065 | 0.264 |
| Diffusion Profiles | 0.091 | 0.347 |
| node2vec + MLP | 0.075 | 0.377 |
| MERW + MLP | **0.115** | **0.501** |
| MERW + RF | 0.079 | 0.261 |
| MERW + 1-NN | 0.081 | 0.308 |
| MERW + 3-NN | 0.093 | 0.333 |
| MERW + 5-NN | 0.085 | 0.331 |

## 5 Conclusion

We have introduced a new way of generating node embeddings using a mass-enhanced random walk approach. For each node in the graph, besides its embedding, we also learn a mass term that captures the influence of this node in the whole network. We demonstrate the efficiency of our algorithm in the drug repurposing task. Our method achieves significant improvement over its competitors in terms of average precision and recall.

